# An integrated database of ion channels

**DOI:** 10.64898/2026.10.12.757514

**Authors:** Kiran Bharat Gaikwad, K. T. Shreya Parthasarathi, Babul Pradhan, John P. George, Jyoti Sharma

## Abstract

Ion channels play critical roles in a wide range of physiological processes, including the propagation of nerve impulses, insulin secretion and regulation of heart rate. Alterations in ion-channels function or expression can contribute to the development of various diseases, including cancers, genetic disorders, and infectious diseases. Ion channels represent the second-largest class of membrane proteins considered potential targets for therapeutic intervention. However, a comprehensive data resource integrating information on human ion channels and their associations with diseases has been limited. Therefore, the Integrated Database of Ion Channels (IDIC) was developed, comprising information on 493 human ion channels. The database contains a broad range of ion-channel annotations, including gene and protein information, structural details, protein signature sequences, protein domains, nuclear localization signals, and signal peptides. It also provides cross-references to several established biological resources, including the Human Genome Organization Gene Nomenclature Committee (HGNC), National Center for Biotechnology Information (NCBI) Gene, Ensembl, and the Protein Data Bank (PDB). Furthermore, IDIC incorporates genomic variations identified in various tumors and disease associations involving ion channels, which have been systematically curated from published literature. Through concise search methods and user-friendly browsing functionalities, IDIC serves as an integrated resource for exploring human ion channels and investigating their structural, functional, genetic, and disease-related characteristics. Thus, the database can facilitate a better understanding of ion channels and their potential involvement in diverse human diseases.

**Database URL:** https://www.idic.co.in/

## Introduction

Ion channels are important signaling molecules that transport ions across biological membranes [2]. The selectivity of ion channels for specific ions, as well as their activation mechanism and kinetics, are highly variable due to their diversity in evolutionary, sequential and structural aspects [5]. This variability across ion channels may be attributed to their involvement in a range of physiological processes including generation of membrane potential, regulation of cell electrical excitability, propagation of action potential along neurons and muscle cells, synaptic release of neurotransmitters, generation of postsynaptic response, and calcium signaling [5]. These structural and functional diversities of ion channels make these proteins imperative. Moreover, ion channels are targets for wide range of endogenous ligands such as hormones and neurotransmitters, and for multiple exogenous toxins and medically important drugs [5].

Alterations in the form of mutations or pathogenic interactions may result in channelopathies due to the involvement of these channels in multiple biological pathways [7-9]. They are key host cell factors for a multitude of viral infections [9]. For survival and replication inside a host, viruses utilize the cellular environment, that is dependent on the flow of ions into and out of the cell [10]. Ca^2+^ channels are involved in viral entry into the cell [11]. Most recently, our group has identified the interactions of ion channels with SARS-CoV-2 proteins and their involvement in insulin secretion, inflammatory mediator regulation of transient receptor potential (TRP) channels, renin secretion, and apelin signaling pathways [12], [13]. Dysregulated potassium channels were identified to be associated with long COVID [14]. Pathogens have evolved to imitate host counterparts in terms of sequence, motif, and structural level in order to utilize host signaling via host-pathogen interactions [15]. Different tumor types including lung adenocarcinoma, basal cell carcinoma, endometrioid carcinoma, acute myeloid leukemia, exhibited mutations in ion channels [16]. Previously, our group has explored their role in tumorigenesis in various cancers [17-22]. *CHRNA4* encodes the α4 subunit of the ligand-gated ion channel nicotinic acetylcholine receptor, was identified in patients with autosomal-dominant nocturnal frontal lobe epilepsy [23]. It has also been reported that the ion channel KCNA3 prevents hepatitis C virus entry during the process of virus-endosome membrane fusion [24]. TRPM2 ion channel is known to function as a global modular of inflammation in neutrophils, by reducing the oxidative response and regulating Ca^2+^ influx and membrane depolarization [25]. Retinal pigment epithelium expresses a wide range of ion channels that are directly or indirectly coupled to retinal pigment epithelium function and may be involved in retinal degeneration as it closely interacts with photoreceptors [26]. Similarly, involvement of deregulated ion channels in 3’,5’-cyclic adenosine monophosphate signaling pathway has been attributed to elevation of cortisol levels in keratoconus patients [27]. Furthermore, dysregulated transient receptor potential cation channels were found to be involved in cellular senescence and peripheral sensitization in Fuchs’ endothelial corneal dystrophy [28]. Growing evidence suggests that bestrophin-1 or calcium voltage-gated channel subunit alpha1 D (CACNA1D) channels contribute to retinal diseases [26].

Considering the importance of ion channels in a variety of metabolic processes, as well as their association in various diseases including cancers, genetic disorders, and infectious diseases, it is crucial to comprehend their role in the disorders. In the current study, 493 ion channels were curated from Human Genome Organization Gene Nomenclature Committee database (HGNC) [29, 30]. Further, the associated data from various publicly available repositories, extensive literature search, and through bioinformatics-based analysis using open-source tools were integrated. The Integrated Database of Ion Channels (IDIC) would be helpful to provide insights into the ion channels that may act as potential targets in several human diseases.

## Methods

### Overview of workflow

Overview of the IDIC is represented in Fig. 1.

**Fig. 1.**
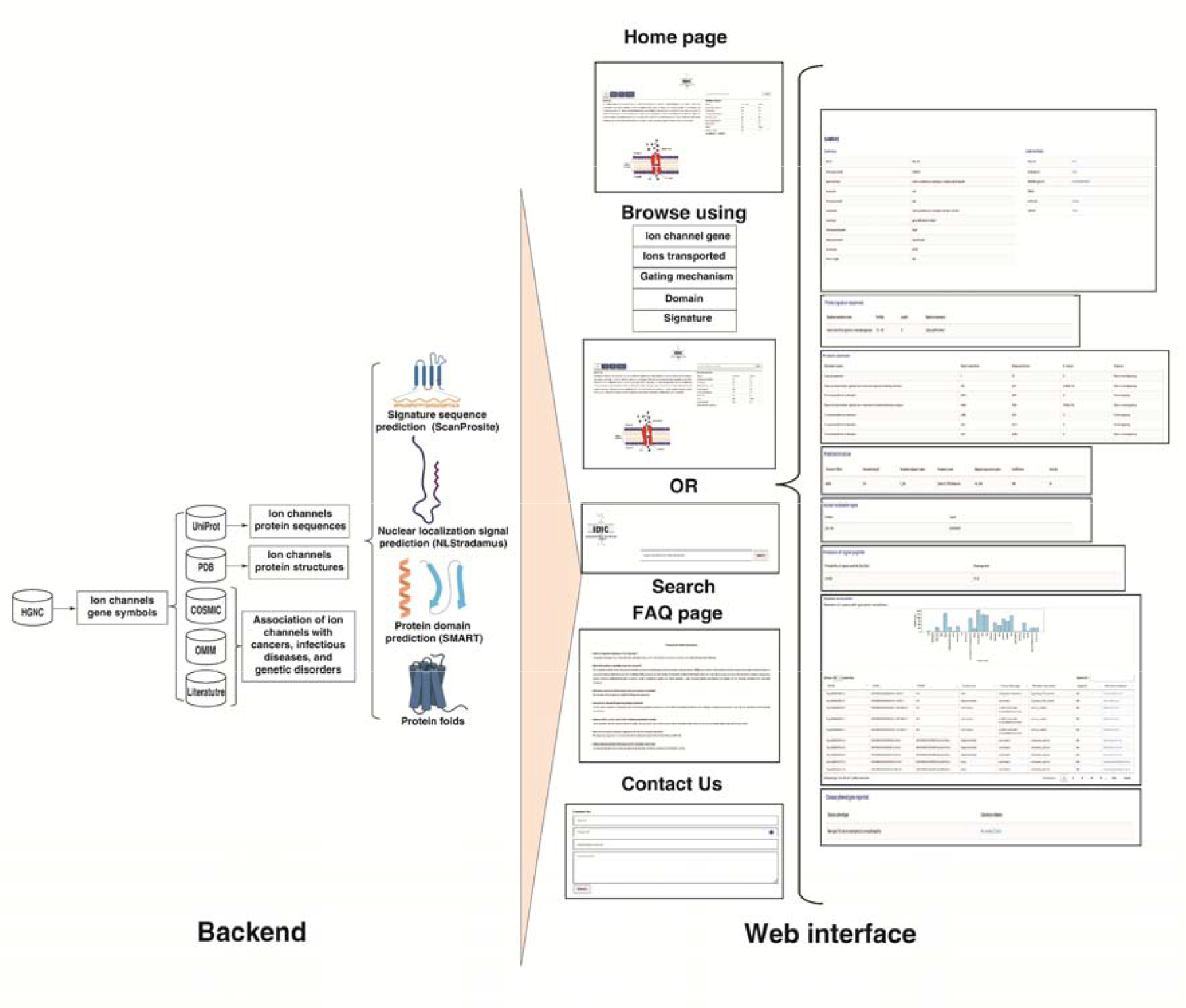
Illustration of the IDIC layout with back-end and front-end.

The data in IDIC was collected from various online resources. It includes information of genes and proteins such as gene symbols, nucleotide sequence length, protein sequence length and details on protein structural features. It also contains information about variations observed in patients with cancer and association of other diseases with ion channels. The database was developed using Django framework with Python (v3.10.11) and MySQL. IDIC offers search option through user input allowing the user to enter the ion channel HGNC symbol to obtain the output including summary, protein features, variations and disease association. The same output can be obtained through browsing options that include searches based on gene symbols, ions transported, gating mechanism, domain name and protein signature sequence name.

### Data collection

1. **Gene information:** The HGNC database was used to collect the ion channel gene symbol and associated gene attributes.
2. **Genomic variations:** The curated ion channels were mapped to the list of annotated variations identified in patients with various tumors from the Catalogue of Somatic Mutations in Cancer (COSMIC) database and extensive literature search.

### Protein attributes

The HGNC gene symbols were used to obtain amino acid sequences of the ion channels from UniProtKB/Swiss-Prot (UniProt Release 2025_02) [31]. These sequences were further used to identify signature sequences using ScanProsite (PROSITR release 2025_02) [32]. Domains were predicted using Simple Modular Architecture Research Tool (SMART) (v10.0) [33-36]. The identifier corresponding to the 3-dimensional (3D) structures for the ion channels were extracted from Protein Data Bank (PDB) [37]. The ion channels were subjected to fold prediction analysis using Protein Homology/analogY Recognition Engine V 2.0 (Phyre2.2) server [36]. The top hit with an identity >= 30% and confidence >= 90% were included in the database. The nuclear localization signals (NLSs) were predicted using NLStradamus (Revision r.9.) [34]. Signal peptides were predicted using SignalP 6.0 [38].

2. **Involvement in diseases:** The detailed information about involvement of various ion channels in other diseases was obtained through literature search.
3. **External links:** IDIC provides the links for external databases including HGNC, National Center for Biotechnology Information, ENSEMBL, PDB, Online Mendelian Inheritance in Man (OMIM) and UniProt [30, 39-43].

### Data search and browse

A user-friendly web interface has been designed for “IDIC” using Django framework. IDIC provides a keyword search option that is a quick search box present in all the Hyper-text markup language (HTML) pages in the web interface. The user is allowed to perform a quick search for any of the 493 ion channels using HGNC symbol of the ion channel. Secondly, 5 search modules – ‘Ion channel’, ‘Ions transported’, ‘Gating mechanism’, ‘Domain’ and ‘Signature’ in Browse page were incorporated to allow user to search the ion channel based on (i) ion channel HGNC symbols, (ii) Ions transported (iii) Gating mechanism, (iv) Domain present in the ion channel and (iv) Protein signature sequence name. The browse by ions transported and gating mechanism option would aid the user in identifying the ion channel when the user is not aware of the HGNC gene symbol. Further the option would allow the user to identify the ion channels that share a similar domain or signature sequence providing insights on the structural similarities between ion channels.

### Database organization and web interface

The data collected were stored and managed in a MySQL relational database management system. It was organized in 13 tables namely, IDIC_main, IDIC_external_links, IDIC_domain, IDIC_signatures, IDIC_nls, IDIC_signal_peptide, idic_predicted_structure, IDIC_variations, IDIC_diseases, and IDIC_ions_transported. The middle layer that would serve as a moderator between the MySQL database and the web interface was developed using Python (v3.10.11) scripts based on Django (v.5.0.3) framework. The web interface of the database was designed using HTML, Cascading Style Sheets and Java script.

## Results

### Information statistics

The information statistics for IDIC data are provided in the Table 1. Information about ion channels and their available/predicted PDB structures is provided in Table 2.

**Table 1.** Information statistics for IDIC. SNV – Single nucleotide Variation.

| **Category** | **Number of features** | **Number of ion channels** |
| --- | --- | --- |
| Protein signature sequences | 62 | 251 |
| Protein domains | 146 | 489 |
| 3-dimensional structures | 122 | 130 |
| Predicted structures | 184 | 357 |
| Nuclear localization signals | 161 | 114 |
| Signal peptides | 84 | 84 |
| Variant | 163047 | 489 |
| Disease phenotypes | 317 | 220 |

**Table 2.** Statistics for structure information of ion channels.

|  |  |
| --- | --- |
| Number of ion channels with folds predicted | 357 |
| Number of ion channels with PDB structures | 130 |
| Number of ion channels with no predicted/PDB structure | 46 |

### Database layout and structure

The database was developed to provide a user-friendly interface to explore ion channels. It consists of an initial Home page, that redirects the user to following pages:

#### 1. Browse page

User can browse IDIC using options such as ion channel, ions transported, gating mechanism, domain, and signature.

#### 2. FAQs

This page contains IDIC related probable queries and their answers that could aid the user to understand the results obtained.

#### 3. Contact Us page

This page enables the user to contact the database developers.

On home page, users can search using either the “Browse” option or the search box provided on the top-right corner of the page. Search box option can be used to enter the ion channel HGNC gene symbol and the user will be redirected to the result page for that particular ion channel. The Browse option provides the user to identify the list of ion channels exhibiting the same feature such as ions transported, gating mechanism, domains or signatures.

### Search results

The result page provides information about that particular ion channel under following sections:

#### 1. Summary

This section consists of IDIC identifier, HGNC gene symbol, approved name, synonyms, previous symbols, group name, locus type, chromosomal location, gating-mechanism of the ion channel, and length of the gene sequence and protein sequence of that ion channel, chromosomal location and amino acid sequence length.

#### 2. External links

It consists of external links to various databases including HGNC, NCBI, ENSEMBL, PDB, UniProt and OMIM.

#### 3. Signature

This section provides the name of the signature sequence/s present in the ion channel along with its position, length, and sequence.

#### 4. Domain

This gives the domain name for the domain present in the ion channel, its start and end position in the protein sequence, E-value of the prediction, and the status that would indicate if the predicted domain is overlapping with another domain or not.

#### 5. Predicted structure

This consists of the PDB id, template and query length and aligned region of the query and template, template name, identity and confidence of the prediction.

#### 6. Nuclear localization signal

This provides the NLSs and their positions in the protein sequences of the ion channels.

#### 7. Presence of signal peptide

This section provides the probability of the prediction of signal peptide in that particular ion channel along with the position of cleavage site.

#### 8. Variations

This section contains the bar graphs for variations indicating number of cases in each of the primary site. It contains the Human Genome Variation Society nomenclatures, primary site, primary histology, mutation description, zygosity, and literature evidence.

#### 9. Association with disease phenotypes

This section provides the diseases associated with the ion channel along with the literature evidence.

## Discussion

### Utility of the database

The purpose behind developing IDIC was to provide a platform that would not only help in recognising the overall properties of ion channels but also assist in identifying their importance in various diseases. In this sense, the database will aid the researchers, as it is the comprehensive resource for the known human ion channels and some of their associated diseases. Moreover, it provides the information about ion channels including their protein sequence and structural features such as signature sequences, domains, NLSs, and three-dimensional structures. Cancer associated variations in ion channels will be helpful in understanding their involvement across different tumors. The visualization of these variations through bar graphs will provide more understanding about the most prevalent tumor associated with the respective ion channel.

Following are the use cases supported by the database:

#### Use case 1: Retrieve the information for an ion channel of interest

This was illustrated in Fig. 2.

**Fig. 2.**
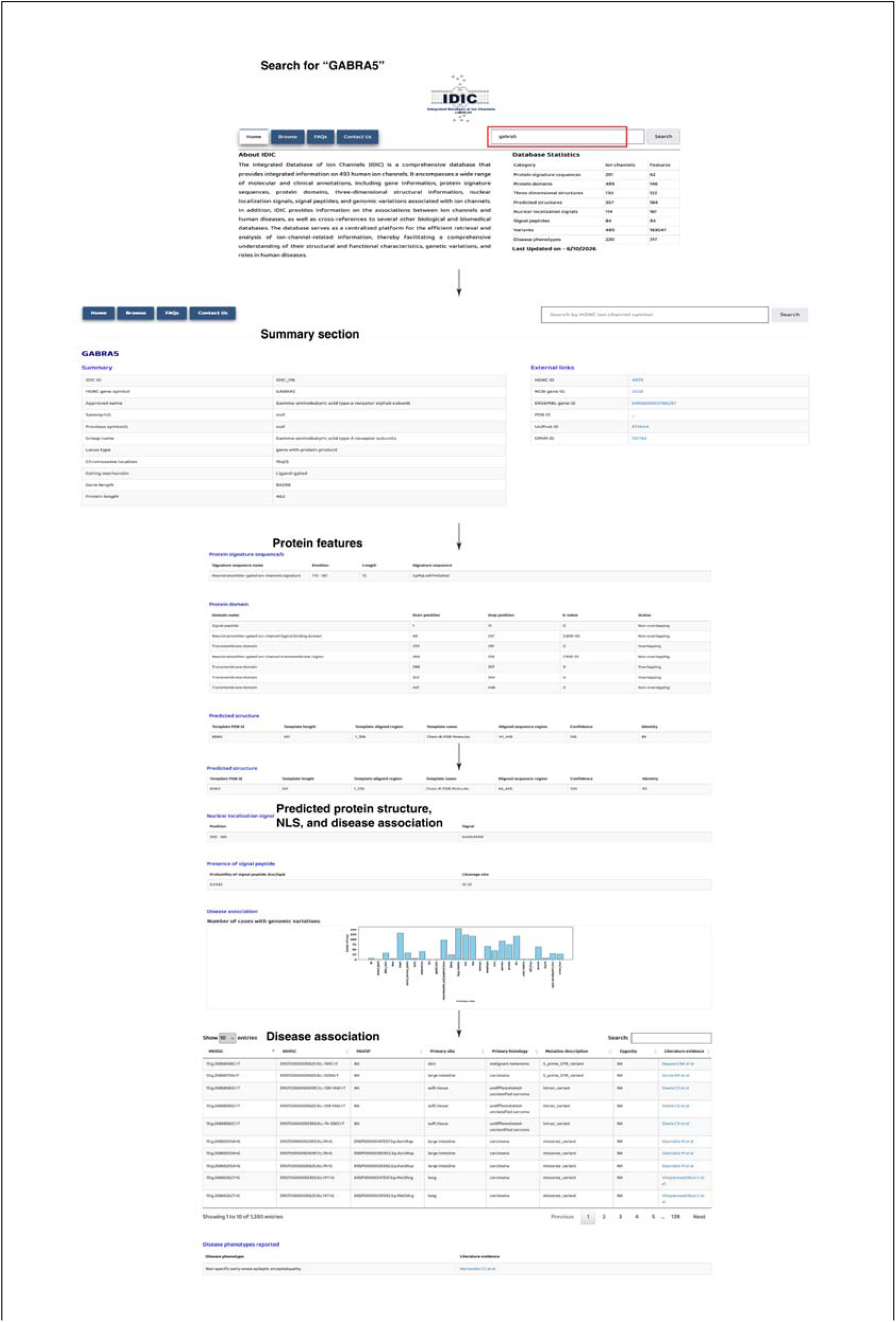

**Fig. 3.**
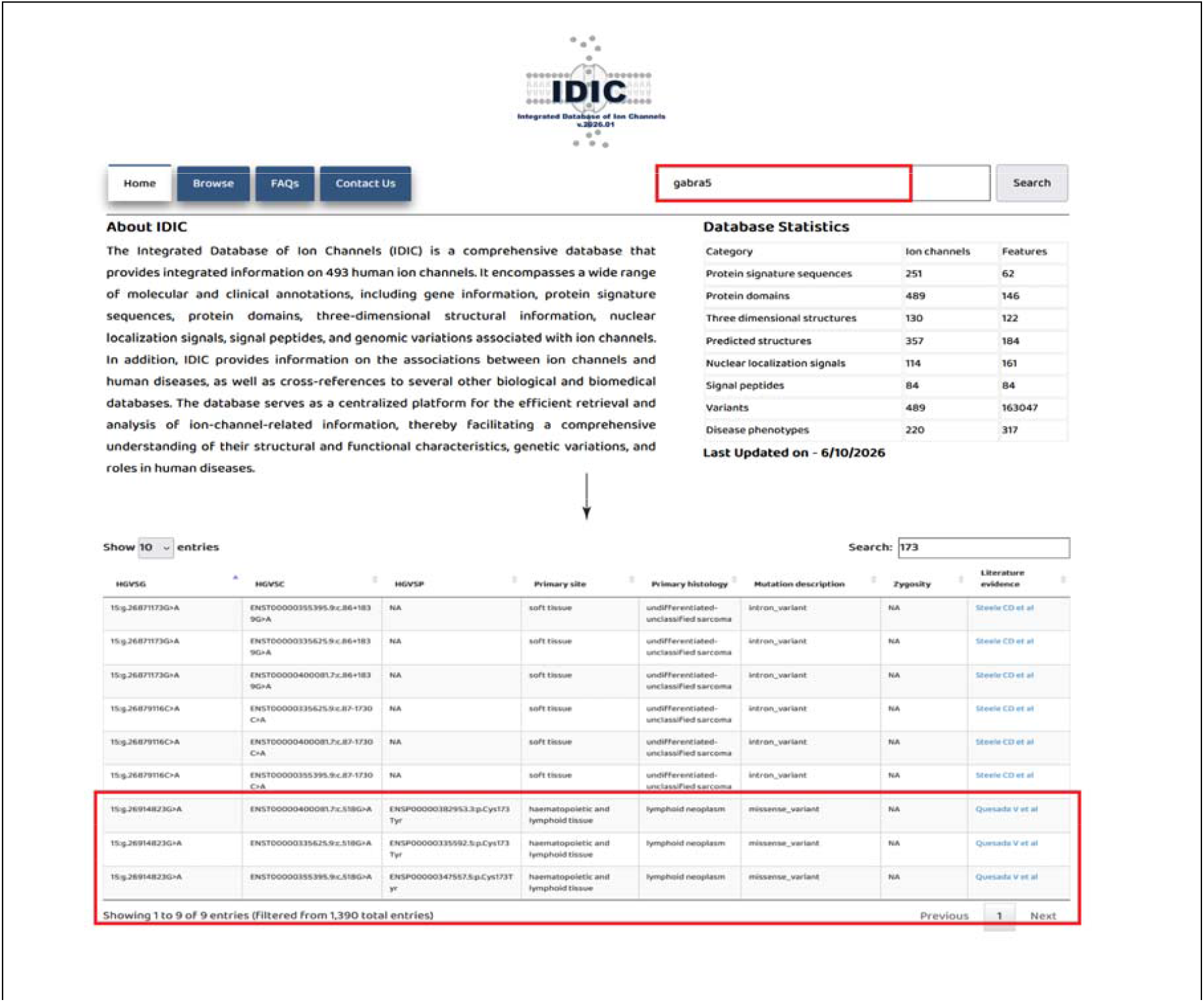
Use case 2 demonstrating how IDIC facilitates linking ion channel variants to functional signatures and disease associations.

User can browse a particular ion channel of interest by inserting the gene symbol in search box. For example, by using “*GABRA5*” as a search term user will get to know about the

##### Summary of GABRA5

HGNC gene symbol, approved name, synonym/s, previous symbol/s, group name, locus type, chromosome location, gating mechanism, gene length, and protein length.

##### External links

HGNC ID, NCBI gene ID, ENSEMBL gene ID, PDB ID, UniProt ID, and OMIM ID.

##### Protein signature sequence/s

Signature sequence name, position, length, signature sequence.

##### Protein domain

Domain name, start position, stop position, E-value, and status.

##### Predicted structure

Template PDB id, Template length, template, aligned region, template name, aligned sequence region, confidence, and identity.

##### Nuclear localisation signal

Position and sequence.

##### Presence of signal peptide

Probability of signal peptide (sec/spi) and cleavage site.

### Disease association

- **Number of cases with genomic variations (for cancers):** This will provide a bar plot indicating the distribution of variants across origine sites of the tumors for *GABRA5*. Current bar plot indicates the frequency of mutations in *GABRA5* is higher in large intestine related cancers followed by breast followed by liver. Additionally, it will provide a paginated table with these columns: HGVSG, HGVSC, HGVSP, primary site, primary histology, mutation description, zygosity, and the literature evidence for the association of the variant with *GABRA5*.
- **Disease phenotypes reported (for genetic and infectious diseases):** This section will provide a table with columns disease phenotype associated with *GABRA5* and literature evidence for the same.

#### Use case 2: Linking ion channel variants to functional signatures and disease associations

1. Search for GABRA5.
2. Navigate to “Disease association” section of the result page.
3. In the search box in top right corner of the table type any position of the signature sequence/domain/NLS/SP to get rows with that number.
4. e.g. After searching for “173” (Start position of the signature sequence in the protein and also present within the predicted domain “Neurotransmitter-gated ion channel ligand binding domain” for which the start position is 49 and end position is 257 with an E-value 2.80E-50), table is filtered based on presence of 173 in the rows and it provides the output with these variants:
  I. ENSP00000382953.3:p.Cys173Tyr
  II. ENSP00000335592.5:p.Cys173Tyr
  III. ENSP00000347557.5:p.Cys173Tyr

This example illustrates the value of integrating multiple levels of ion channel information in IDIC. In particular, the association of a variant with a predicted functional signature provides additional biological context for interpreting its potential relevance, while the linked disease annotation connects the molecular feature to a human disease phenotype. Such integrated information can facilitate the identification and prioritization of variants and ion channel proteins for further functional investigation.

#### Use case 3: Identification of variants in the NLS

Mutations in NLS can contribute to the abnormal trafficking of proteins to the nucleus [44]. This makes it important to verify the presence of literature supported mutations in this region. Following are the steps to identify mutations in the NLS of *GABRA5* and associated cancers (Fig. 4):

**Fig. 4.**
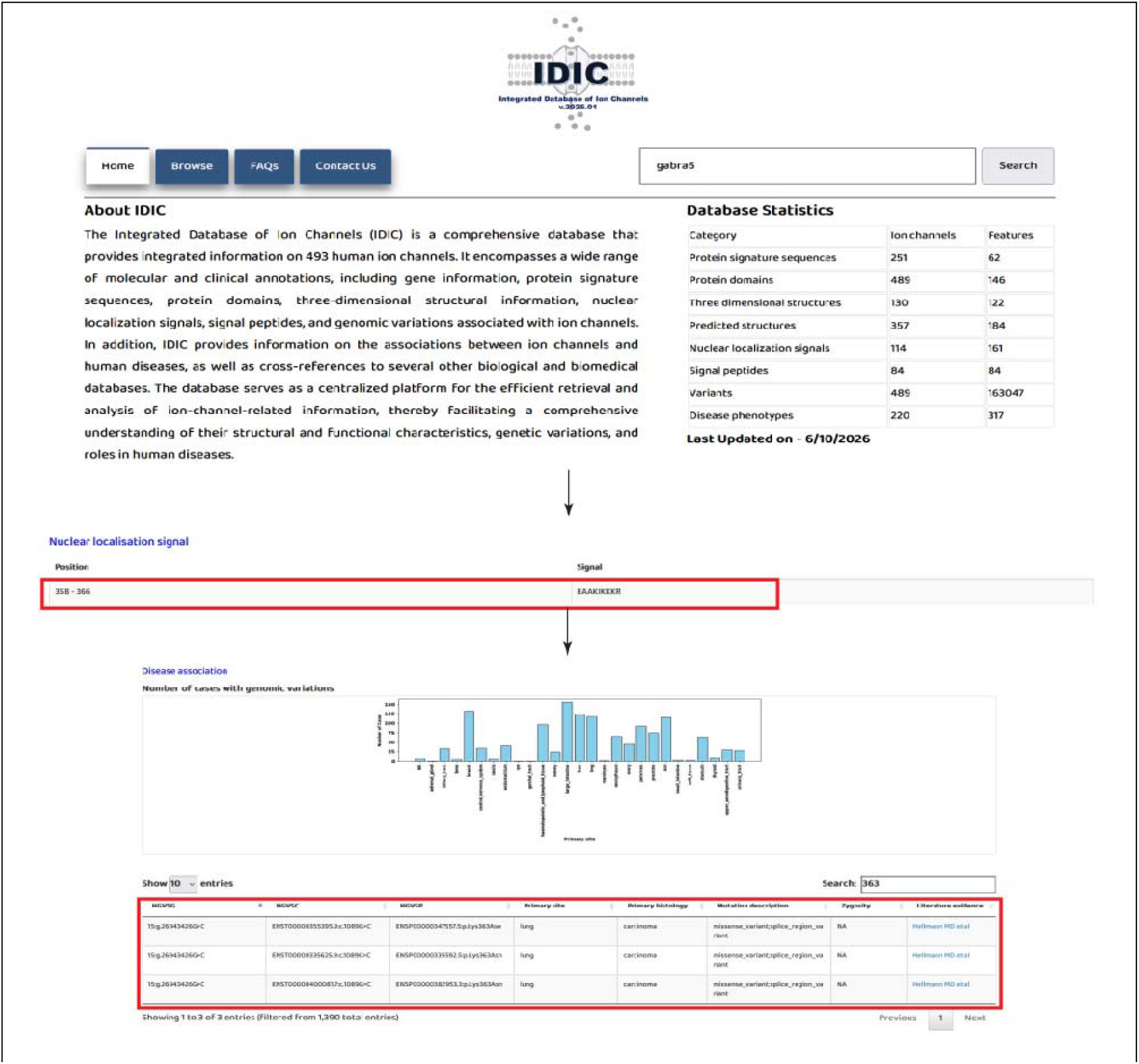
Use case 3 highlighting identification of variants within the nuclear localization signal (NLS) of GABRA5.

1. Search for “GABRA5” using the search box.
2. Na
3. vigate to “Nuclear localisation signal” section.
4. The signal position in the protein is 358-366.
5. Navigate to “Disease association” section and check for presence of mutation at any of this position.
6. Mutations with following HGVSP will be listed:
  I. ENSP00000347557.5:p.Lys363Asn
  II. ENSP00000335592.5:p.Lys363Asn
  III. ENSP00000382953.3:p.Lys363Asn

These mutations are mainly reported in the lung carcinoma [45].

Overall, the IDIC database would serve as a platform for information on ion channels, providing a wide range of annotations, giving more insights about ion channels and their association with diseases (Table 3).

**Table 3.** Comparison of IDIC with other existing ion channel databases.

| Comparison criterion | IDIC | IUPHAR-DB | Channelpedia | ChannelsDB 2.0 | MemProtMD |
| --- | --- | --- | --- | --- | --- |
| Primary objective | One stop solution to obtain all the information about the ion channels. | IUPHAR-DB integrates peer-reviewed pharmacological, chemical, genetic, functional, and anatomical information. | Channelpedia enables collaborative efforts for ion channel information from the scientific literature. | To provide structural information about the position, geometry, and physicochemical properties of ion channels- tunnels and pores. | To provide a computational resource for understanding how integral membrane proteins interact with lipid bilayers. |
| Human ion channels coverage | This resource includes information about 493 ion channels. | The database contains 212 ion channels including 71 ligand-gated and 141 voltage-gated ion channels. | This resource contains 187 ion channels. | The database contains about 350 proteins with predicted transmembrane pores. | 3,506 membrane-protein structures/simulations; number of ion channels not specified. |
| Gene/protein information | Gene- Yes<br>Protein- Yes | Gene- Yes<br>Protein- Yes | Gene- Yes<br>Protein- Yes | Gene- No<br>Protein- Yes | Gene- No<br>Protein- Yes |
| Structural information | Yes | Available through cross linking to PDB. | Yes | Yes | Yes |
| Disease association for cancer and genetic/infectious diseases separately | Yes | Disease information and association is available through cross referencing to other databases. Clear distinction/separate section is not available for cancer and infectious/genetic diseases. | Clear distinction/separate section is not available for cancer and infectious/genetic diseases. | No | No |
| Genomic variants | Yes | Yes | Yes | Yes (Available through UniProt mutation annotation data) | No |
| Literature evidence | Yes | Yes | Yes | Yes | Yes (Associated with source structures/publications) |
| Cross-database identifiers | Yes | Yes | Yes | Yes | Yes (Includes cross-references for UniProt and PDB) |
| Main biological use | Comprehensive analysis of human ion channels and their disease associations, enabling disease-focused exploration, channel-disease relationship mapping, and identification of candidate ion channels for research purpose. | It can be used in pharmacological profiling of ion channel targets and their ligand/drug interactions. | Integrated analysis and computational modeling of ion channel function, with emphasis on neuronal electrophysiology. | Structural analysis of protein channels—including tunnels and pores—and characterization of molecular and ion transport pathways. | Understanding membrane-protein structure and function in a lipid-bilayer environment through molecular-dynamics simulation and analysis of protein-lipid interactions. |
| References |  | [1] | [3] | [4] | [6] |

### Uniqueness and importance

The IDIC database is the first online platform that provides comprehensive information for 493 human ion channels and their association with various diseases.

Several other resources of ion channels including IUPHAR-DB, Channelpedia, ChannelsDB, and MemProtMD are available[1], [3], [4], [6]. However, these resources do not provide information regarding all the 493 known ion channels. Comparison of these databases with IDIC is provided in Table 3. IDIC is a centralized repository for the information of the human ion channels and their associations across disorders.

The added value of IDIC is primarily its integrated, human ion channel, and disease-oriented framework. The database is flexible to update and reformulate. Specifically:

1. **Data types:** IDIC integrates information on ion channels together with their associated disease information and related biological annotations in a unified resource. In contrast, the existing resources are generally specialized in particular aspects of ion channel biology. For example, ChannelsDB emphasizes structural characterization of channels, pores and tunnels; Channelpedia emphasizes ion channel information and electrophysiological/computational models; IUPHAR/BPS GtoPdb primarily focuses on pharmacological targets and ligand interactions; and MemProtMD focuses on membrane-protein molecular-dynamics simulations and protein–lipid interactions.
2. **Curation and integration**: IDIC was designed to systematically provide information for human ion channels (at the level of genes and proteins) and their disease associations, allowing channel-related information to be interpreted in a disease context. This disease-oriented integration distinguishes IDIC from resources whose primary curation objectives are structural, pharmacological, electrophysiological, computational-modeling, or membrane-simulation data.
3. **Search and retrieval capabilities:** IDIC provides an integrated framework for retrieving ion channel information together with disease-associated information, thereby allowing users to navigate from ion channels to their associated diseases. This facilitates cross-linking of channel and disease information without requiring users to manually combine information from multiple specialized resources.
4. **Biological use cases:** IDIC is intended to support disease-oriented research related to human ion channels, including searching for of channel–disease relationships, interpretation of ion channel biology in human disease, and generation of hypotheses for further investigation. Thus, IDIC complements specialized resources such as ChannelsDB, Channelpedia, MemProtMD, and IUPHAR/BPS GtoPdb.

It is easier to include further information as and when available without disturbing the current database layout. Simple and user-friendly interface of the database will allow wider access to the information for both experimentalists as well as bioinformaticians.

## Future updates

The database will be updated in future as and when new information will be available on the ion channels. Besides updating currently available information, the database will be frequently updated for additional information and features. This will also be done based on the users’ feedback.

Additional features that are planned to be implemented include addition of protein-protein interactions. This would include graphical representation of the biological interactions of ion channels among themselves and with other proteins. In near future, the database will offer more diseases in terms of variations.

## Limitations

As with any biological database, IDIC has certain limitations that we aim to address through future improvements and updates. One potential limitation is the continuity and accuracy of the boundaries of the protein domains predicted for ion channels. Since these domains were identified using computational prediction methods, their use in further research may require experimental validation to confirm their precise boundaries and functional relevance. Another limitation concerns the availability of structural information. In the current version of IDIC, structure prediction was performed using sequence identity and confidence cut-offs. Consequently, predicted or experimentally determined structures are not available for 46 ion channels. This information will be updated in future versions as additional structures become available.

## Acknowledgements

J.S. would like to thank the Council of Scientific and Industrial Research (CSIR), Government of India [37WS(0114)] and Indian Council of Medical Research (ICMR), Government of India [FIW-2025-01-00001091] for the research support. J.P.G. is supported by the CSIR [37WS(0114)], K.B.G. is supported by the ICMR [FIW-2025-01-00001091], and B.P. is recipient of the Junior Research Fellowship [DBT/2025-26/IOB/2976] from the Department of Biotechnology (DBT), Government of India.

## Author’s contribution

J.S., K.B.G., K.T.S.P.: Conceptualization and Methodology, J.S.: Supervision, K.B.G., K.T.S.P., B.P., J.P.G.: Data Curation, K.B.G., K.T.S.P.: Software, Visualization, K.B.G., K.T.S.P.: Writing—original draft preparation, K.B.G., K.T.S.P., B.P., J.P.G.: Writing—review and editing. All the authors read and approved the final manuscript.

## Data availability

The resource is available at https://www.idic.co.in/

